# A combined RNA-seq and whole genome sequencing approach for identification of non-coding pathogenic variants in single families

**DOI:** 10.1101/766717

**Authors:** Revital Bronstein, Elizabeth E. Capowski, Sudeep Mehrotra, Alex D. Jansen, Daniel Navarro-Gomez, Mathew Maher, Emily Place, Riccardo Sangermano, Kinga M. Bujakowska, David M. Gamm, Eric A. Pierce

## Abstract

Inherited retinal degenerations (IRDs) are at the focus of current genetic therapeutic advancements. For a genetic treatment such as gene therapy to be successful an accurate genetic diagnostic is required. Genetic diagnostics relies on the assessment of the probability that a given DNA variant is pathogenic. Non-coding variants present a unique challenge for such assessments as compared to coding variants. For one, non-coding variants are present at much higher number in the genome than coding variants. In addition, our understanding of the rules that govern the non-coding regions of the genome is less complete than our understanding of the coding regions. Methods that allow for both the identification of candidate non-coding pathogenic variants and their functional validation may help overcome these caveats allowing for a greater number of patients to benefit from advancements in genetic therapeutics. We present here an unbiased approach combining whole genome sequencing (WGS) with patient induced pluripotent stem cell (iPSC) derived retinal organoids (ROs) transcriptome analysis. With this approach we identified and functionally validated a novel pathogenic non-coding variant in a small family with a previously unresolved genetic diagnosis.

## Introduction

Inherited retinal degenerations (IRDs) are a leading cause of blindness, altogether affecting >2million people worldwide. IRDs are characterized by progressive degeneration of photoreceptor and/or retinal pigment epithelial (RPE) cells of the retina with variable age of onset and rates of degeneration^1^. Despite tremendous ongoing efforts and research into various therapeutics, treatments for IRDs remain limited. Currently there are two regulatory agency-approved treatment approaches, retinal prosthesis implants and gene augmentation therapy for IRD caused by mutations in the *RPE65* gene ^1–7^. The eye is a prime candidate for gene therapy approaches due to its relative immune-privilege, surgical accessibility and ease of non-invasive monitoring. In addition, IRDs are Mendelian disorders caused by mutations in single genes in the vast majority of cases. Owing to these favorable circumstances several gene/genetic therapy clinical trials have been initiated for IRDs, including those caused by mutations in the *ABCA4, CEP290, CHM, CNGA3, CNGB3, MYO7A, RPGR, RS1,* and *USH2A* ^6^.

As each genetic therapy targets a specific gene, for a patient to be considered for treatment they must obtain a reliable genetic diagnosis. Difficulties inherent to genetic diagnostics are rooted in the fact that every individual carries millions of DNA variants in their genome ^8, 9^. The large majority of the DNA variants are found in non-coding regions of the genome such as intergenic and intronic regions. Since non-coding sequences can better tolerate sequence variation compared to coding sequences, most of these variants are benign and do not lead to disease. Still, some non-coding variants are found to be pathogenic by altering gene expression and/or splicing patterns ^10–12^. As a result, non-coding variants are among the hardest to classify and thus under-diagnosed ^13, 14^. Algorithms exist that predict the effect of a non-coding variant on gene expression or splicing based on analysis of the DNA sequence alone, but their accuracy for diagnostic purposes remains undetermined ^15–18^.

In order to functionally test the effects of non-coding variants one needs to quantify the level of gene expression and analyze the splicing patterns of the presumably affected genes. When multiple variants in multiple genes need to be evaluated, advanced methods for whole transcriptome analysis are advantageous. Indeed, previous studies successfully utilized large RNA-seq datasets from tissue biopsies to identify novel non-coding pathogenic variants ^19, 20^. For example, Evrony *et al* narrowed down a linkage analysis in a very large pedigree to a single non-coding variant using RNA-seq. This non-coding variant was shown to cause intron retention in the *DONSON* gene and is most likely the genetic cause of microcephaly-micromelia syndrome (MMS) in this population ^19^. Cummings *et al* used 184 skeletal muscle RNA-seq samples available through Genotype-Tissue Expression resource (GTEx) ^21^ as a reference panel for 50 patients with undiagnosed muscle disorders. This comparison led to a genetic diagnosis for 17 previously unsolved families and identification of several splice altering variants ^20^.

Such studies depend on the availability of large, publicly available RNA-seq datasets and/or on a large cohort of patients. They also require biopsy samples from a clinically relevant tissue or cell type. Both gene expression and splicing are tissue-specific ^22, 23^ owing to restricted availability of transcription and splicing factors with variable usage of regulatory DNA sequences ^24–27^. Consequently, DNA variants in such regulatory sequences can have tissue-specific outcomes on gene expression and splicing ^18, 28^. Thus, analyzing RNA from a clinically relevant tissue or cell type is crucial to obtain a more focused and reliable diagnostic result. When a clinically relevant tissue is not accessible, *ex vivo* surrogate models can sometimes suffice. Indeed, a study aimed at discerning the genetic causality of patients with monogenetic neuromuscular disorders found that t-myotubes, skeletal myotubes derived by myoD overexpression in fibroblasts, accurately reflected the muscle transcriptome and faithfully revealed pathogenic variants ^29^.

In this study, we aimed to develop a pipeline that would detect putative non-coding pathogenic mutations in a small family. We present a pilot study performed in a five member family with two siblings affected by cone dysfunction syndrome in which we successfully identify and functionally validate a novel deep intronic variant without the use of large reference datasets. Obtaining a clinically relevant tissue from IRD patients is not possible as the retina cannot be safely biopsied. To overcome this limitation, we have made use of patient-derived induced pluripotent stem cells (iPSCs) that were differentiated *in vitro* to form retinal organoids (ROs)^30^. ROs have been shown to recapitulate many aspects of human retinal structure and function ^30, 31, 32–39, 40^. We show that the RO transcriptome is much closer to the transcriptome of normal human retina than other more readily available diagnostic tissues. More importantly, we show for the first time that analysis of a patient-derived RO transcriptome can successfully detect pathogenic deep intronic variants that activate cryptic splice sites, leading to a new genetic diagnosis. Our approach can lead to a larger number of patients to be eligible for genetic therapies.

## Materials and Methods

### Human Subjects

The study was approved by the institutional review board at the Massachusetts Eye and Ear (Human Studies Committee MEE in USA) and adhered to the Declaration of Helsinki. Informed consent was obtained from all individuals on whom genetic testing and further molecular evaluations were performed.

### Pluripotent Stem Cell Induction

Tissue samples were obtained with written informed consent in adherence with the Declaration of Helsinki and with approval from institutional review boards at the University of Wisconsin-Madison and Massachusetts Eye and Ear Infirmary. Blood samples from 4 individuals from family OGI-081 (197, 198, 200 and 340) were collected and reprogrammed by Cellular Dynamics, Inc (now FUJIFILM Cellular Dynamics, Inc) as custom MyCell products. Three independent clones from each individual were karyotypically normal, expressed pluripotency markers and successfully differentiated to retinal organoids (^41^: lines 1579, 1580, 1581 and 1582). Stem cells were maintained on Matrigel (ThermoFisher) in either mTeSR1 (WiCell) or Stemflex (ThermoFisher) and passaged with either Versene or ReLeSR (STEMCELL Technologies).

### Retinal Organoid (RO) Differentiation

Differentiation of iPSCs was performed as previously described ^41^. Briefly, embryoid bodies (EB) were lifted with either 2 mg/ml dispase or ReLeSR and weaned into Neural Induction Media (NIM: DMEM:F12 1:1, 1% N2 supplement, 1x MEM nonessential amino acids (MEM NEAA), 1x GlutaMAX and 2 mg/ml heparin (Sigma)) over the course of 4 days. On day 6, 1.5 nM BMP4 (R&D Systems) was added to fresh NIM and on day 7, EBs were plated on Matrigel at a density of 200 EBs per well of a 6-well plate. Half the media was replaced with fresh NIM on days 9, 12 and 15 to gradually dilute the BMP4 and on day16, the media was changed to Retinal Differentiation Media (RDM: DMEM:F12 3:1, 2% B27 supplement, MEM NEAA, 1X antibiotic, anti-mycotic and 1x GlutaMAX). On days 25-30, optic vesicle-like structures were manually dissected and maintained as free floating organoids in poly HEMA (Sigma)-coated flasks with twice weekly feeding of 3D-RDM (RDM + 5% FBS (WiCell), 100 μM taurine (Sigma) and 1:1000 chemically defined lipid supplement) to which 1 μM all-trans retinoic acid (Sigma) was added until d100. Live cultures were imaged on a Nikon Ts2-FL equipped with a DS-fi3 camera.

### Immunocytochemistry and Microscopy

Organoids were fixed in 4% paraformaldehyde at room temperature for 40 min, cryopreserved in 15% sucrose followed by equilibration in 30% sucrose, and sectioned on a cryostat. Sections were blocked for 1 hr at room temperature (RT) in 10% normal donkey serum, 5% BSA, 1% fish gelatin and 0.5% Triton then incubated overnight at 4°C with primary antibodies diluted in block. Table S1 lists primary antibodies, dilutions and sources. Slides were incubated with species-specific fluorophore-conjugated secondary antibodies diluted 1:500 in block, for 30 minutes in the dark at RT (Alexa Fluor 488, AF546 and AF647). Sections were imaged on a Nikon A1R-HD laser scanning confocal microscope (Nikon Corporation, Tokyo, Japan).

### DNA Sequencing

DNA was extracted from venous blood using the DNeasy Blood and Tissue Kit (Qiagen, Hilden, Germany). OGI-081-197 underwent GEDi sequencing as described previously ^42^. All five family members underwent whole exome and PCR-free whole genome sequencing. Sequencing was done at the Genomics Core at Massachussets Eye and Ear as described previously ^43^.

### RNA Sequencing

For transcriptome analysis, ROs from at least 2 different clones per individual were harvested at approximately day160 (early stage 3), lysed in 350 µl buffer RLT+ME from the RNeasy mini kit (Qiagen), snap frozen on dry ice and stored at −80C. At a later time samples were defrosted on ice and passed through QIAshredder columns (Qiagen). Subsequently, Total RNA was extracted per the manufacturer’s instructions. RNA quality and quantity was assessed on an Agilent 2100 Bioanalyzer, RIN number ranged between 9.6-9.9. For each sample 1μg of total RNA spiked with 1.2ng Sequins (v2) controls ^44^ was used to generate RNA-seq paired-end libraries with the Illumina TruSeq Stranded Total RNA kit. Ribosomal RNA was removed with the Ribo-Zero Human/Mouse/Rat kit. Libraries were multiplexed and sequenced on an Illumina HiSeq 2500 instrument for 101 cycles.

### Bioinformatics

Whole exome sequence data was analyzed in house ^43^ and whole genome data was analyzed in collaboration with the Broad Institute of MIT and Harvard using methodology described previously ^20^. Briefly, BWA was used for alignment. GATK was used for single nucleotide polymorphism and insertion/deletion calls. Additional variant annotation was performed using the Variant Effect Predictor (VEP) ^45^. Variants of interest were limited to polymorphisms with less than 0.005 allelic frequency in the gnomAD and ExAC databases ^8, 9^. Whole genome copy number analysis, with consideration of structural changes, was done using Genome STRiP 2.0 ^46^.

For analyses of RNA-seq data, read quality was assessed with FastQC v0.11.3 (Babraham Bioinformatics, Cambridge, UK) and MultiQC v1.2 ^47^. Reads were aligned to the human genome version GRCh37 by the STAR v2.5.3a ^48^ aligner in two-pass mode within the sample and across replicates for each sample sets. Annotations were derived from the Human GENCODE v19 (Ensemble74).

FeatureCount v1.5.2 ^49^ from the Subread package, was used to generate gene expression matrix with the following non-default settings, reads must be paired, both the pairs must be mapped, use only uniquely mapped reads, multi-mapped reads are not counted, chimeric reads are not counted and strand specificity turned on. Anaquin ^50^ was further used to evaluate alignment sensitivity and gene expression. Here sensitivity indicates the fraction of annotated regions covered by alignments of the reads by STAR (Table S2). No limit of quantification or limit of detection was reported.

For discovery of novel or known alternative splicing events we used a combination of CASH v2.2.1 ^51^ and MAJIQ v1.1.3a ^52^. CASH was operated with default settings. MAJIQ was run with a minimum of 5 reads for junction detection and 10 reads for the calculation of delta percent spliced-in (dPSI). EdgeR (v3.2.2) ^53^ was used to perform differential gene expression with default settings. Data normalization was performed using trimmed mean of M-values (TMM). Next, the raw read counts were converted to transcript per million (TPM) expression values. The Picard tools v1.87 and RSeQC v2.6.4 ^54^ were used to calculate mean fragment length. The approach implemented in Kallisto ^55^ was used to covert raw reads to TPM values. An average TPM of the third lowest Sequins between test and control samples was calculated and used as cutoff. TPM values for the GTEx samples used in figure3 were downloaded from the GTEx portal. The human normal retina (HNR) samples ^56^ and the ENCODE skin samples were reanalyzed as described above. The ENCODE skin samples were used for the analysis performed with MAJIQ as they were generated with a stranded total RNA-seq library same as our RO samples. In contrast the GTEx samples were generated with an mRNA non-stranded library.

### RT-PCR and cloning

RT-PCR was conducted using SuperScript IV first-Strand synthesis system (ThermoFisher Scientific, Waltham, MA). Exon 14b was amplified with primers F: GACATGTTGCTAAGATTGAAATCCGT from exon 14 and R: GACCCAGCTTTCAGAGTAACCAGAAC from exon 15 using Phusion polymerase (NEB, Ipswich, MA). The longer band containing exon14b was then excised from the gel and purified using Zymoclean gel DNA recovery kit (Zymoresearch, Irvine, CA) and cloned using pGEM-T Easy Vector System (Promega, Madison, WI). The plasmid was used in a transformation into Subcloning Efficiency DH5α Competent Cells (Invitrogen, Carlsbad, CA). The plasmid was isolated with Zyppy Plasmid Miniprep Kit (Zymoresearch, Irvine, CA). All procedures described in this section were conducted according to the manufacturer instructions. The DNA sequence of Exon 14b was found to be: GCCAGGTGCAGTGGCTCACGACTGTAATTCCAACACTTTGGGAGGCCAAGG TGGCAGGATCACATAAGTCCAGGAGTTCAAGACAAGCCTGGACAACATG.

## Results

### Unresolved genetic analysis of family OGI-081

The pilot study reported here was conducted on a five member family with two siblings shown by clinical testing to be affected by a cone dysfunction syndrome (Figure1a). Both affected patients had nystagmus and decreased vision from infancy and at age 8, OGI-081-197 was also noted to have photophobia. Visual acuity for both affected patients measured 20/150-200 and remained stable for 3 years. Full field electroretinogram (ffERG) testing of retinal function for OGI-081-197 was significant for reduced and delayed cone photoreceptor responses, with normal rod photoreceptor response amplitudes. Optical coherence tomography (OCT) imaging of the retina showed retinal degenerative changes in the fovea (Figure1b). In addition to their retinal disease, both affected patients were found to have Chiari malformations. Interestingly, vision phenotypes such as photophobia, vision loss and nystagmus have been reported as accompanying symptoms in some forms of Chiari malformations^57–60^. Unfortunately, despite evidence that Chiari malformations have a hereditable component ^61–63^, the genes involved are not yet well defined ^59, 64, 65^. For that reason we could not rule out the possibility that the vision phenotypes and Chiari malformations share a common genetic causality.

**Figure 1:**
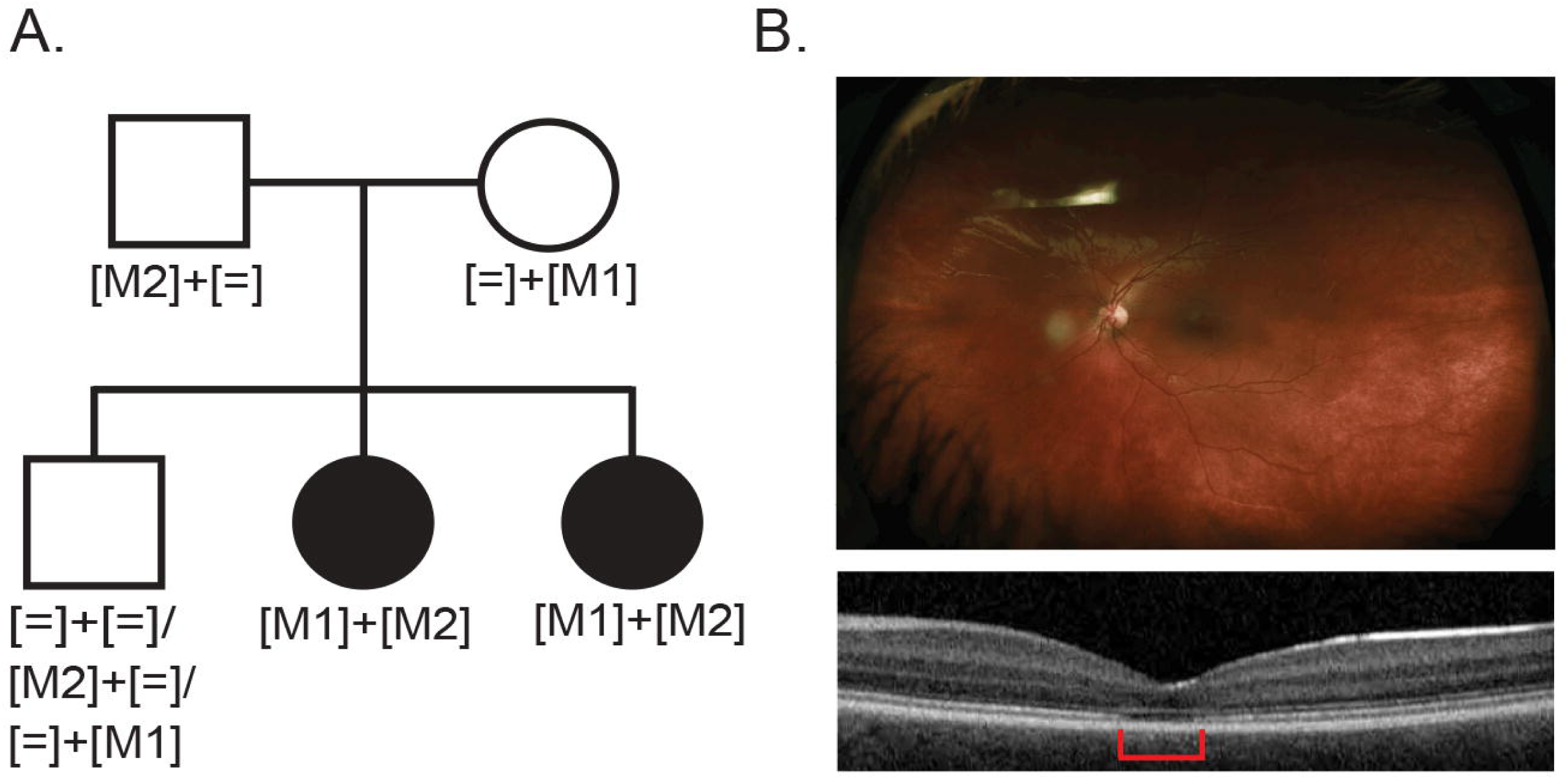
Family OGI-081 variant segregation scheme and retinal phenotypes. A. The OGI-081 pedigree with variant segregation scheme. B. Fundus (upper panel) and OCT image (lower panel) for the OGI-081-197 at age 8, area of retinal degeneration is indicated by the red bar.

The phenotypes segregation in OGI-081 is indicative of a recessive mode of inheritance. Thus, for genetic testing we searched for genes that have putatively pathogenic variants in both alleles (Figure1a). Selective exon capture based genetic diagnostic testing was performed using the Genetic Eye Disease (GEDi) test ^42^. Since this did not identify a clear cause of disease, whole exome sequencing (WES) for the five members of the family was performed. Both the GEDi testing and WES identified a single rare variant in the *CNGB3* gene (c.1148delC, p.Thr383IlefsTer13) which has been reported to be pathogenic ^66–68^, but a second rare variant in *CNGB3* was not identified, nor were other potential causative genetic variants forthcoming for the two affected members of the family. *CNGB3* mutations are among the most common causes of cone dysfunction syndrome, but to the best of our knowledge, Chiari malformations have not been reported as an accompanying symptom in *CNGB3* patients ^69, 70^. Moreover, the 1.75e^-3^ gnomAD allele frequency of the p.Thr383IlefsTer13 variant is higher than expected for recessive pathogenic variants and two homozygous individuals are reported in the gnomAD database. Since it has been proposed that up to 1 in 4-5 individuals in the general population may be a carrier of null mutations in IRD genes ^71^ it was possible that the presence of variant p.Thr383IlefsTer13 was an incidental finding.

We therefore decided to test for two possible disease scenarios. One is that *CNGB3* accounts for the cone dysfunction syndrome and the second allele is a non-coding variant. In this case the Chiari malformation has a separate, unrelated causality. The second possibility is that a novel disease-causing gene is responsible for both the cone dysfunction syndrome and the Chiari malformation. To test these hypotheses, we performed both whole genome sequencing (WGS) and RNA-seq analysis of a surrogate retinal tissue to determine whether the combination of these orthogonal investigations could yield a clear genetic solution.

Analysis of DNA variants detected by WGS identified 3268 segregating rare variants that could be sorted into 8191 allelic pairs in 642 genes (TableS3). The variant ranked at the top of the list of potential causes of disease remained the known pathogenic variant c.1148delC; p.Thr383IlefsTer13 in the *CNGB3* gene. However, a second coding variant once again was not found in this gene. Next, we set to establish a clinically relevant surrogate transcriptome for the human retina.

### The iPSC-derived retinal organoid (RO) transcriptome can be used as a surrogate for a human retinal biopsy

Patient-derived iPSCs were generated from peripheral blood monocytes of all members of family OGI-081 excluding OGI-081-199. The iPSCs were subjected to an *in vitro* differentiation process to generate ROs with attached RPE (Figure 2a) ^30^. RPE was specifically retained in the ROs so as to concurrently identify potential mutations in RPE genes as well as photoreceptor genes (Figure2b). The ROs were kept in culture for 160 days (early stage 3 ^30^), a time point at which outer segments are visible by light microscopy and cone and rod cells are clearly distinguished by immunocytochemistry (Figure2b, c-h). At da160, ROs were harvested for total RNA isolation, library preparation, and RNA-seq analysis (Table1).

**Figure 2:**
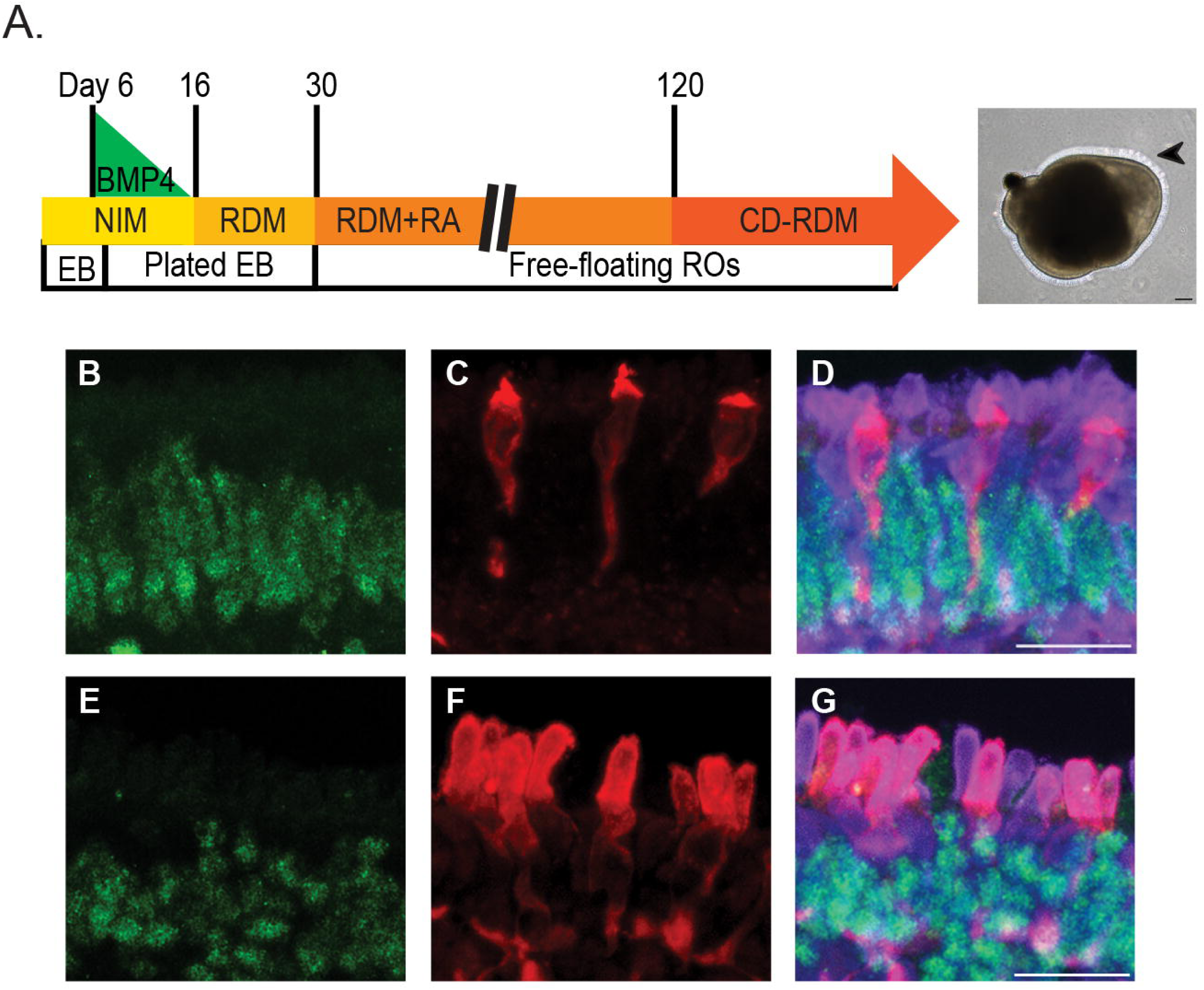
RO differentiation. A. Schema of the differentiation process and a light microscopy image of a typical RO. Arrow head indicating photoreceptors, scale bar = 100 microns. B-G. Immunocytochemistry on cryosections of ROs. B&E. NR2E3 staining of rod nuclei (Green). C&F. Mature cones show staining of cone opsins in the cone photoreceptor outer segments (Red). C. S opsin. F. ML opsin. D&G. Overlay of rod and cone staining. All cones are stained with ARR3 (Purple). Scale bars = 20 microns.

**Table 1:** Sample level alignment report and QC summary. A high average unique percentage alignment rate is reported.

| SampleName | RawReads<sup>a</sup> | HQReads<sup>b</sup> | % UniqAligned<sup>c</sup> |
| --- | --- | --- | --- |
| OGI-081-197-1 | 159,976,749 | 129,671,633 | 87.73 |
| OGI-081-197-2 | 256,399,765 | 234,701,223 | 90.52 |
| OGI-081-197-3 | 166,237,226 | 155,442,261 | 88.88 |
| OGI-081-198-1 | 167,327,080 | 145,058,230 | 88.66 |
| OGI-081-198-2 | 197,037,935 | 179,026,100 | 89.13 |
| OGI-081-198-3 | 191,485,904 | 177,663,416 | 90.15 |
| OGI-081-340-1 | 175,199,604 | 156,566,678 | 91.01 |
| OGI-081-340-2 | 227,928,431 | 216,526,830 | 89.53 |
| OGI-081-340-3 | 137,096,260 | 129,192,437 | 90.47 |
| OGI-081-340-4 | 151,572,341 | 140,982,711 | 90.59 |
| OGI-081-340-5 | 164,908,758 | 154,550,198 | 99.66 |

<sup>a</sup> Raw reads count (RawReads), <sup>b</sup> Filtered high quality reads (HQReads), <sup>c</sup>
Percent of uniquely aligned reads (% UniqAligned).

In order to evaluate if day 160 ROs can be used as informative surrogates for the adult human retinal transcriptome, we examined the expression levels of 270 known IRD genes reported in the RetNet database. For this analysis, we compared an in-house dataset composed of 3 post-mortem human normal retinas (HNR, N=3)^56^ to ROs derived from the unaffected sibling (N=5) or skin or whole blood samples taken from the GTEx database (Figure 3 and Table S4)^21^. Skin and blood represent tissues that are more readily available – and thus commonly used – for surrogate diagnostic testing. We found 224 IRD genes to be expressed in HNR (TPM>1). Interestingly we were able to detect expression of 254 IRD disease genes in the RO samples (Figure 3 and TableS4). The higher detection rate of disease-causing genes in ROs compared to HNR is most likely because of overall higher TPM values in RO samples (Figure 3a&b), possibly due to higher RNA quality compared to post-mortem HNR samples. In addition, the presence of RPE and photoreceptors with varied maturation statuses in the RO samples could be contributing factors to this finding. As expected, skin and blood expressed lower numbers of IRD genes (188 & 130 respectively) at much lower TPM values (Figure 3a&b and Table S4). Since IRD genes were very poorly represented in the blood samples, we excluded blood from further analysis.

**Figure 3:**
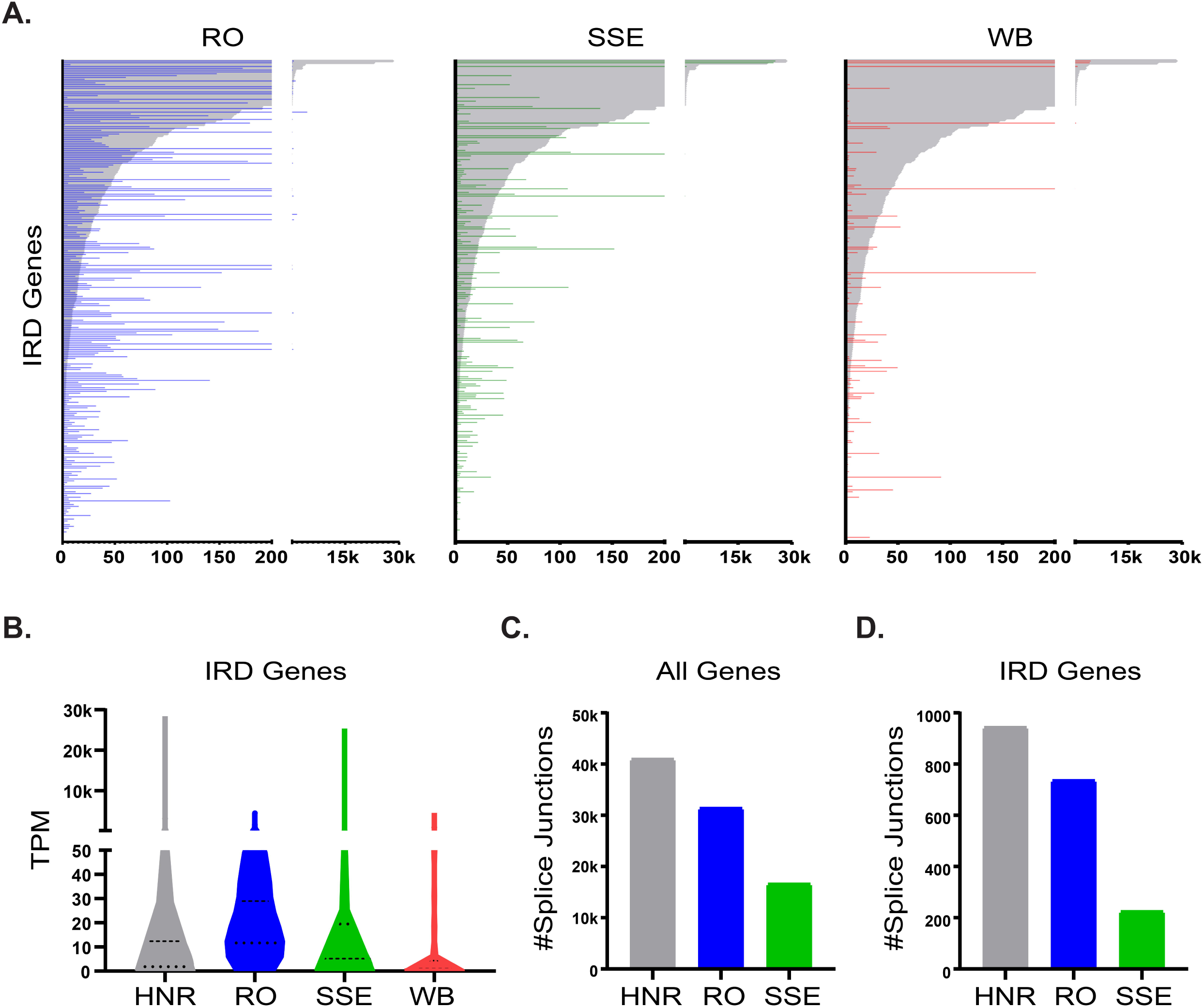
Comparison of IRD gene expression and splice junctions. Human Normal Retina (HNR; N=3, gray), RO from the unaffected sibling (N=5, blue), Skin-Sun Exposed (SSE; N=473, green) and Whole Blood (WB; N=407, red). A&B Average TPM values of IRD genes. A. IRD genes are sorted by their expression in HNR overlaid with RO, SSE or WB. B. Violin plot. C&D Number of splice junctions detected by MAJIQ. C. All annotated genes. D. IRD genes.

We next examined the complexity of the HNR, RO and skin transcriptomes, which is reflected by the multitude of isoforms that are produced from each gene locus ^72, 73^. Isoform diversity that occurs via alternative splicing of the pre-mRNA can be represented by the splice junctions detected in each gene locus. We used the MAJIQ ^52^ algorithm to detect splice junctions in HNR and RO samples of the OGI-081 unaffected sibling, and the ENCODE database ^74, 75^ to detect splice junctions in corresponding skin samples. The skin samples from the ENCODE database were more suited for this analysis then the GTEx samples due to RNA-seq library type (see Materials and Methods). We found a comparable number of splice junctions in the HNR and RO samples (41,121 and 31,535 (76%) respectively) but a much lower number in the skin samples (16,713 (40%)) (Figure 3c and TableS5). This result is probably due to the fact that HNR and ROs are composed of a more diverse cell population than skin. More importantly, in IRD genes, the gap in complexity between the HNR (946 junctions) and ROs (739 (78%)) as compared to skin (227(24%)) is even greater (Figure 3d & Table S5). Thus, the RO transcriptome provided a close facsimile of the human retinal and IRD transcriptomes at the gene expression and splicing pattern levels, whereas the skin transcriptome did not.

### Detection of a novel non-coding pathogenic variant in *CNGB3*

In order to find the underlying genetic cause of the retinal degeneration in OGI-081 affected patients, we conducted differential splicing and gene expression analyses of the RNA-seq data obtained from day 160 ROs of affected versus unaffected siblings. The differential splicing analysis was conducted with CASH ^51^ and MAJIQ ^52^ algorithms, and we used edgeR algorithm ^53^ for differential gene expression. CASH detected 106 differential splicing events in 101 genes (Table S6), while MAJIQ detected 522 differential splicing junctions in 260 genes (Table S7). A comparison to the 642 genes with DNA variant pairs indicated that only two genes, *CNGB3* and *NCALD*, had altered splicing patterns and a DNA variant in each of their alleles that segregated according to disease status in OGI-081 (Figure 4a).

**Figure 4:**
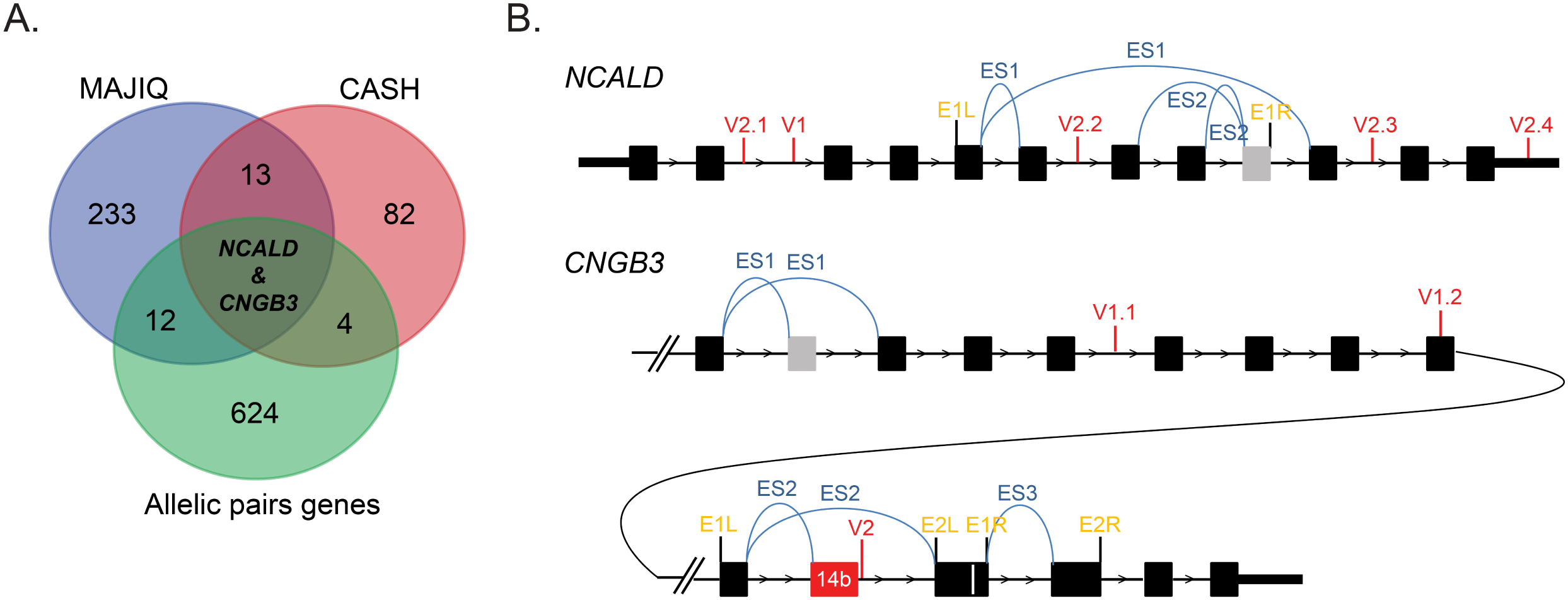
Alternative splicing in the *NCALD* and *CNGB3* genes. A. Venn diagram of genes found to have alternative splicing events in OGI-081 comparison of affected vs. unaffected siblings and genes found to have segregating allelic pairs (green). Alternative splicing analysis was conducted by MAJIQ (blue) and CASH (red). B. Collapsed diagram of exons (black boxes) from all isoforms of the *NCALD* and *CNGB3* genes. DNA variants (red); MAJIQ (blue) and CASH (yellow) alternative splicing events (E). Events detected by MAJIQ are depicted as split reads arches. The event range detected by CASH is depicted by the left (L) and right (R) borders. Genomic locations of variants, junctions and event borders are given in table S6.

For the *NCALD* gene, the alternative splicing events identified by CASH and MAJIQ were isoform switching events between minor isoforms (data not shown) whose biological significances are unclear. In addition, the DNA variants in this gene are located 15-50kb away from the nearest alternative junction (Figure 4b and Table S8). Given such large distances, it is not likely that these variants could cause the alternative splicing events. Moreover, the differential gene expression analysis did not find the *NCALD* gene to be differentially expressed between the affected and unaffected siblings (Table S9). Taken together, these results indicate that the *NCALD* gene variants are most likely not the genetic cause for the disease phenotypes observed in OGI-081.

We next examined the *CNGB3* gene locus. Based on segregation in the family, we found one allele carrying the intronic variant chr8:g.87618576G>A while the second allele carried the known pathogenic variant c.1148delC; p.Thr383IlefsTer13, and a second intronic variant chr8:g.87676221T>C (Table S3). Both MAJIQ and CASH detected an alternative splicing event spanning variant chr8:g.87618576G>A. In contrast, no alternative splicing events were found to span variant chr8:g.87676221T>C (Figure4b and TableS6-8). Close examination of the alternative splicing event spanning variant chr8:g.87618576G>A revealed that it incorporated a cryptic exon into *CNGB3* in RNA samples taken from both affected siblings but not from the unaffected sibling (Figure 5a&b). The cryptic exon is spliced between canonical exon 14 and exon 15 and will therefore be termed exon14b from hereon. The inclusion of exon 14b was validated by RT-PCR and subsequent cloning and Sanger sequencing of the novel longer isoform from the two affected siblings (Figure 5b).

**Figure 5:**
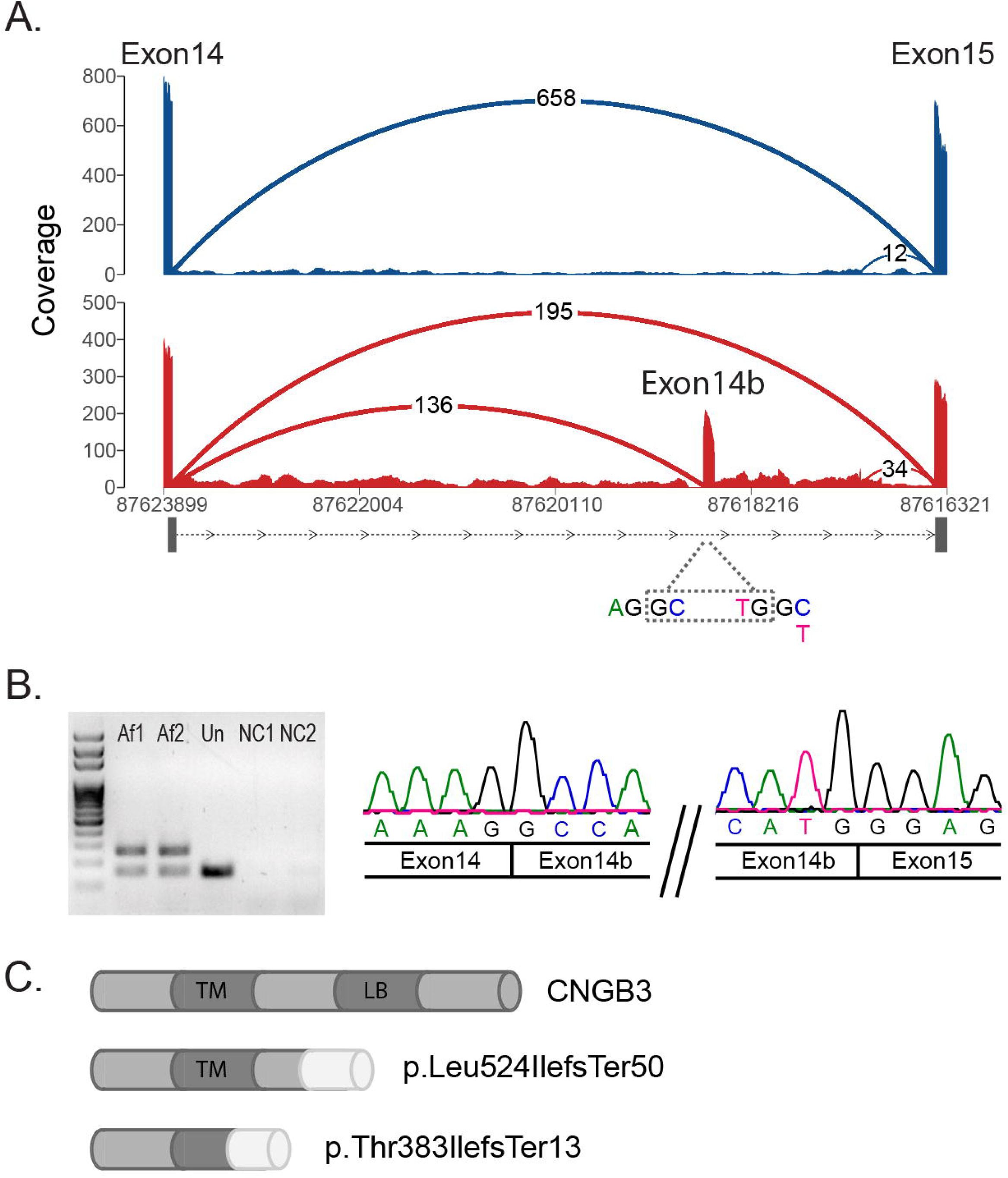
Aberrant splicing of *CNGB3* in the affected vs. unaffected siblings. A. Sashimi plot presenting RNA-seq results showing a cryptic exon spliced into the isoform as a result of the intronic variant chr8:g.87618576G>A. The cryptic exon is only present in the affected sibling (lower panel, red) and not in the unaffected sibling (upper panel, blue). The splice junction between exon 14b and exon 15 is not represented by split reads in the Sashimi plot due to an alignment error (FigureS1). B. RT-PCR using primers from the canonical exon14 and exon 15. All three siblings express the normal size isoform lacking exon 14b (lower band). A larger abnormal band containing exon14b (upper band) is present in the two affected siblings Af1 and Af2 but not in the unaffected sibling (Un). Negative controls lacking RNA template in the RT reaction (NC1) and NC1 used as template for PCR amplification (NC2). Sanger sequencing of the larger band confirming the inclusion of exon 14b. C. Schematic representation of the protein domains in W.T CNGB3 and the two mutant alleles found in the affected siblings of OGI-081.

Both the addition of exon 14b to the *CNGB3* transcript as a result of variant chr8:g.87618576G>A, and the single base pair deletion in the second allele carrying variant c.1148delC; p.Thr383IlefsTer13, lead to a frame shift and subsequent premature termination. We therefore expected both alleles to undergo nonsense mediated decay (NMD) with down regulation of *CNGB3* mRNA levels in the affected siblings as compared to the unaffected sibling. However, contrary to our expectations, *CNGB3* expression was not significantly down regulated in our RNA-seq dataset, as analyzed using the edgeR program. A comparison of each of the affected siblings with the unaffected sibling yielded log_2_FC values of −1.05 and −1.13, indicating slightly lower expression levels in the affected siblings that did not reach statistical significance (p-values of 0.13 and 0.16 respectively (Table S9)). These two frame shift alleles are predicted to encode truncated proteins. The protein encoded by the exon 14b including isoform is predicted to maintaining a full transmembrane domain but lack the ligand binding domain of CNGB3 (Figure 5c). Similarly, the protein encoded by the isoform carrying the known pathogenic variant c.1148delC; p.Thr383IlefsTer13 is predicted to have a truncated transmembrane domain in addition to lacking the ligand binding domain (Figure 5c). In order to determine whether the truncated CNGB3 proteins are being translated, we performed immunohistochemistry on ROs from the exon 14b allele carrier parent (OGI-081-200) and one affected sibling (OGI-081-197, Figure 6). CNGB3 is a subunit of the cone cyclic nucleotide-gated (CNG) channel, which localizes to cone photoreceptor outer segments in chicken and mice ^76, 77^. We have also validated the localization of human CNGB3 to cone photoreceptor outer segments in the human retina (Figure S2). We therefore immunostained ROs for CNGB3 and ML opsin, the latter serving as a marker for photoreceptor outer segments. For these studies, stage 3 ROs were kept in culture for a total of 262 days, allowing cones full opportunity to mature and localize ML opsin and CNGB3 to the photoreceptor outer segments. As expected, CNGB3 co-localized with ML opsin in cone photoreceptor outer segments in the parent (Figure 6c), with weaker staining observed in inner segments as well (Figure 6b&c), presumably due to mislocalization of truncated CNGB3 protein produced by the exon 14b including allele. In ROs from the affected sibling, where both alleles are predicted to result in truncated proteins, CNGB3 was only observed diffusely in the cell body and in inner segments; i.e., no co-localization with ML opsin was observed in cone photoreceptor outer segments (Figure 6e&f). Taken together, results from the differential splicing analysis indicate that the likely cause for the inherited retinal degeneration in OGI-081 is two pathogenic alleles in *CNGB3* - the known pathogenic allele p.Thr383IlefsTer13 and the novel deep intronic allele chr8:g.87618576G>A; NM_019098.3:c.1663 – 2137C>T; pLeu524 IlefsTer50.

**Figure 6:**
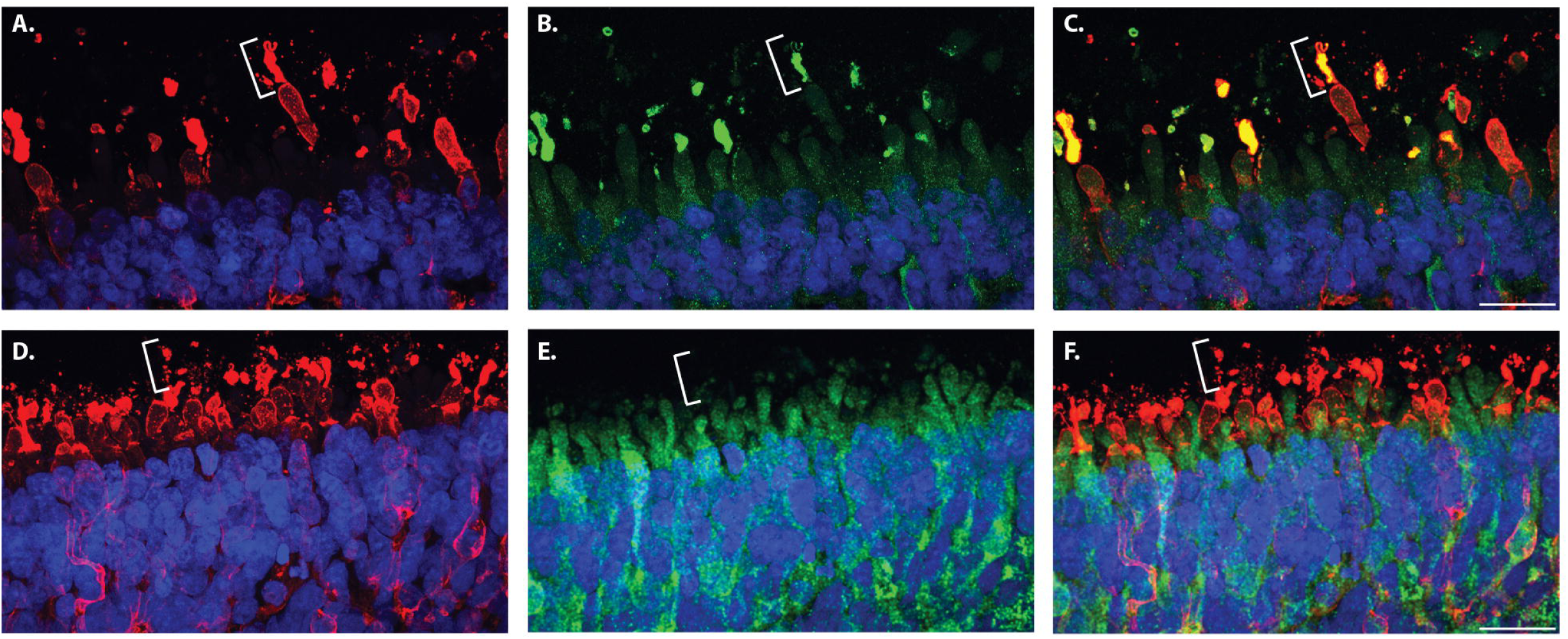
Mislocalization of the CNGB3 truncated proteins. Immunocytochemical analysis of day 262 ROs from the heterozygous parent OGI-081-200 (A-C) and an affected sibling OGI-081-197 (D-F). In the heterozygote, both ML opsins (red) and CNGB3 (green) are localized to the photoreceptor outer segments whereas in the affected sibling, CNGB3 localizes to the photoreceptor inner segments. An exemplary photoreceptor outer segment is indicated by the white brackets. Nuclei are counterstained with DAPI (blue). Scale bars = 20 micron

### Splicing prediction algorithms

With the identification of the non-coding pathogenic variant in *CNGB3,* we set out to examine the mechanism by which it promotes the inclusion of exon 14b. We analyzed the splice junctions surrounding exon14b with the variant analysis tool Alamut Visual. Alamut Visual incorporates three splicing predictors capable of analyzing deep intronic variants, SpliceSiteFinder-like (SSF)^78^, MaxEntScan ^79^ and NNSPLICE^80^. All three algorithms predicted chr8:g.87618576G>A to strengthen a cryptic donor splice site (DSS) (Table 2). All three algorithms also detected a potential acceptor splice site (ASS) at position c.1663-2238, exactly where our Sanger sequencing indicated the acceptor site of exon 14b resides. Interestingly, exon 14b ASS is a stronger than the one located at exon 15 (Table 2). It is plausible that the availability of this acceptor site and its ability to compete with the acceptor site of exon 15 contributed to the effect of variant chr8:g.87618576G>A on the splicing pattern of *CNGB3* in the affected siblings. In addition we noticed the presence of a second even stronger alternative ASS 54bp upstream of exon 15 ASS. It is possible that this secondary competitor further weakens the exon 15 ASS thus enhancing the effects of variant chr8:g.87618576G>A.

**Table 2:**
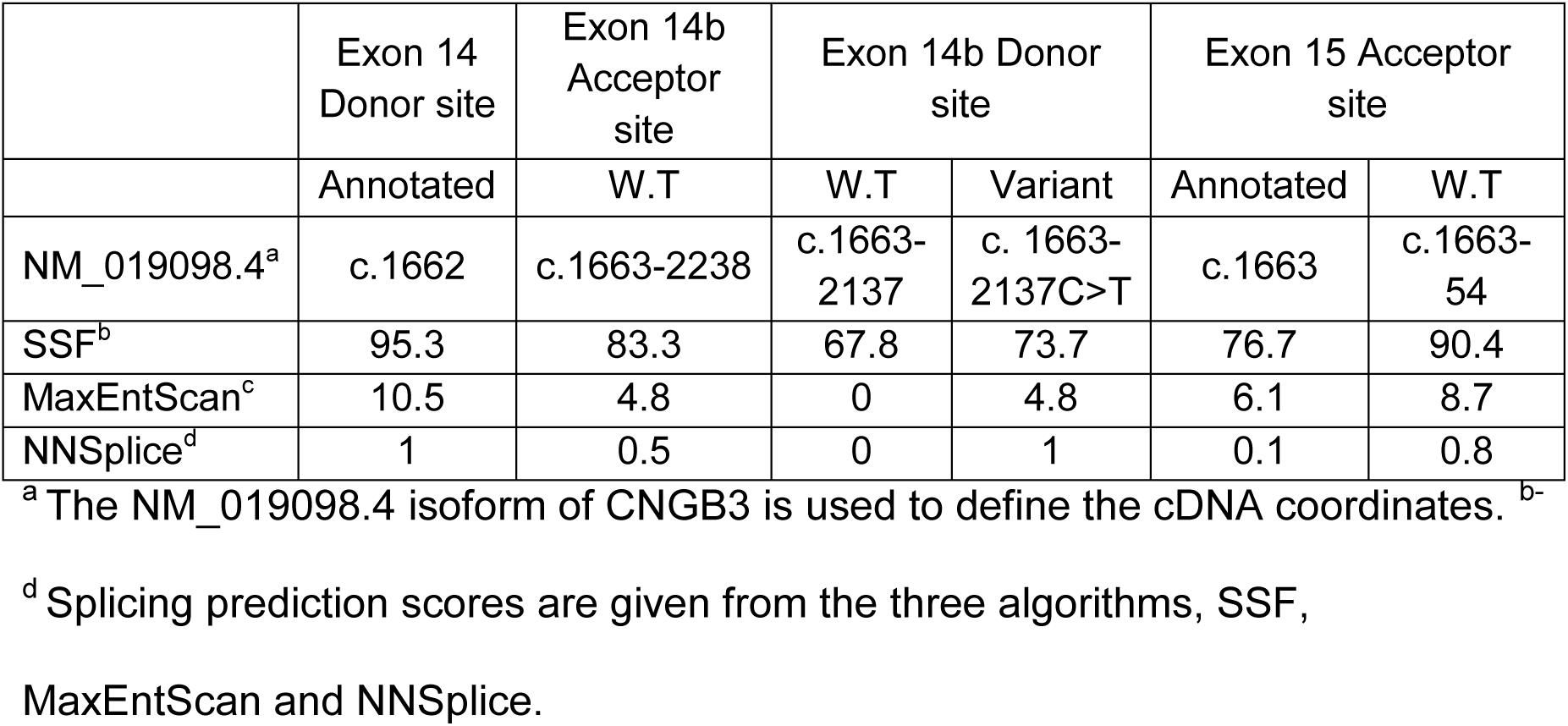
Splicing junctions involved with the inclusion of exon14b.

|  | Exon 14 Donor site | Exon 14b Acceptor site | Exon 14b Donor site |  | Exon 15 Acceptor site |  |
| --- | --- | --- | --- | --- | --- | --- |
|  | Annotated | W.T | W.T | Variant | Annotated | W.T |
| NM_019098.4<sup>a</sup> | c.1662 | c.1663-2238 | c.1663-2137 | c.1663-2137C>T | c.1663 | c.1663-54 |
| SSF<sup>b</sup> | 95.3 | 83.3 | 67.8 | 73.7 | 76.7 | 90.4 |
| MaxEntScan<sup>c</sup> | 10.5 | 4.8 | 0 | 4.8 | 6.1 | 8.7 |
| NNSplice<sup>d</sup> | 1 | 0.5 | 0 | 1 | 0.1 | 0.8 |

<sup>a</sup> The NM\_019098.4 isoform of CNGB3 is used to define the cDNA coordinates. <sup>b-</sup>
<sup>d</sup> Splicing prediction scores are given from the three algorithms, SSF,
MaxEntScan and NNSplice.

Next, using OGI-081 as a true positive case, we tested whether splicing prediction algorithms could be used to prioritize candidate non-coding splicing altering variants, circumventing the need for RNA-seq analysis. We annotated the 3,268 rare variants with allelic pairs identified in OGI-081 with two splicing prediction programs. (i). Alamut Batch that makes its prediction by the combined calculations of the same splicing predictions algorithms as Alamut visual but is capable of calculating the effects of multiple variants. (ii). SpliceAI, a deep neural network tool, to predict splice junctions from pre-mRNA transcript sequence ^18^. Alamut batch calculated a high probability for altering splicing for 532 variants in 315 genes (Table S10). Although variant chr8:g.87618576G>A, the novel pathogenic variant identified in this study, was predicted by Alamut Batch to strongly activate a cryptic donor site the large number of additional candidate variants make this tool too cumbersome for identification of candidate non-coding pathogenic variants. For SpliceAI, to identify synonymous exonic, near intronic, and deep intronic variants predicted to affect splicing at a validation rate of 40% the authors used Δ Score greater than or equal to 0.2, 0.2, and 0.5 respectively. Out of the variants segregating in OGI-081only eight had scores 0.2<0.5 and only one, variant Chr9:g.86536129C>T, received a Δ Score >0.5 (Table S11). Variant chr8:g.87618576G>A, the novel pathogenic variant identified in this study as activating a cryptic donor splice site, was calculated by SpliceAI to have a donor gain Δ Score of 0.3 well below the 0.5 cutoff for deep intronic variants. Thus, had we used SpliceAI splicing predictions as a filter to identify potential causal variants for functional validations, variant chr8:g.87618576G>A would have been overlooked. The ASS of exon 14b was not identified by either algorithm and therefore could not have been used to highlight variant chr8:g.87618576G>A as a more plausible pathogenic variant.

## Discussion

We present here an unbiased approach based on the combination of WGS and RNA-seq data to identify and functionally validate pathogenic non-coding variants without the use of large datasets. We show that *ex vivo* models, such as iPSC derived ROs, can serve as a surrogate source of a patient’s own retinal tissue for RNA and protein analyses. IRDs are currently at the focus of gene therapy advances and several clinical trials are underway, including a trial for *CNGB3* gene augmentation therapy ^6^. This work was aimed at expanding the number of patients eligible for clinical trials and forthcoming therapies. Indeed, our findings here make the two affected siblings of OGI-081 eligible to participate in ongoing clinical trials for *CNGB3* gene therapy. Our approach is applicable to any inherited disease, both WGS and RNA-seq techniques are commercially available, gold standards are being established and the analysis tools are readily accessible ^47, 48, 49–55, 81, 82^. *Ex vivo* organoid models are being developed for a multitude of tissues including brain ^83, 84^, kidney ^85^, liver ^86, 87^ and lung ^88^.

Non-coding variants present a challenge for a correct genetic diagnosis that is imperative for a successful genetic therapy. The combination of WGS and RNA-seq methodologies allows us to both detect non-coding variants and evaluate their functionality throughout the genome. Indeed, a similar approach has already been successfully employed to diagnose diseases where RNA could be harvested from biopsies of disease-relevant tissue ^19, 20^. These studies relied on the availability of large control datasets of RNA-seq samples from unaffected individuals and/or a large cohort of patients ^19, 20^. Our work shows that the correct diagnosis of non-coding variants is possible without reliance on such resources. WGS analysis of all five members of OGI-081 and segregation analysis of the variants within the family narrowed down the search from tens of thousands of variants to a few hundred with allelic pairs. We then used RNA-seq analysis comparing two affected siblings to an unaffected one as an orthogonal approach to identify genes with altered splicing or expression in disease. Thus identifying the deep intronic variant chr8:g.87618576G>A as a novel pathogenic variant in the *CNGB3* gene. The iPSC derived ROs served both as a source of disease relevant transcriptome and as a system for functional validation of the truncated proteins. In future studies, for families with a single patient the parents may serve as control samples, so that each parent controls for the effect of the allele inherited from the other parent making our approach applicable even for ultra-rare diseases.

Once the deep intronic variant was detected and validated we were able to use that prior knowledge to identify additional factors that may have contributed to the inclusion of exon 14b such as the availability of the cryptic acceptor site of exon 14b and the comparative weakness of the exon 15 acceptor site. Such complex dependencies are a prime example as to why sequence based predictions of splicing patterns are hard to compute. Still, several splicing predictors in the Alamut Visual software were able to detect the increase in the splicing probability of the cryptic donor site as a result of variant chr8:g.87618576G>A. This prompted us to test whether such splicing predictors can be used as preliminary filters to identify candidate pathogenic variants for “gene by gene” validation methods, circumventing the need for RNA-seq analysis of ROs. We found that the more established approach represented by the Alamut Batch method of combining the calculations of several splicing predictors that are designed to identify known splicing motifs yielded too many candidates for gene by gene validation. In contrast, the more recent approach of deep neuronal networks algorithms, represented by SpliceAI, failed to assign high probability to the true positive variant chr8:g.87618576G>A. Still, in cases where some prior knowledge can help prioritize variants or highlight ones with lower than expected scores these methods may yet be helpful. In cases where no prior knowledge can help prioritize candidate variants, such as in patients where both pathogenic alleles are non-coding, and especially in cases where a cell type relevant for functional validation is not available, the approach established here is preferable.

## Appendices

## Differential gene expression analysis

As mentioned above, differential gene expression analysis was less informative in the OGI-081 datasets. We compared gene expression levels from each of the affected siblings to that of the unaffected sibling (Table S9). We found 401 genes to be consistently down regulated, of which 27 were Y linked as expected given that the two affected siblings are females while the unaffected sibling is a male. We excluded these Y linked genes from further analysis. Tools are not currently available to filter out non-Y linked genes that may be differentially expressed between the sexes under normal conditions in ROs. Of the remaining 374 down regulated genes, 29 also contained allelic variant pairs (Table S3). In addition, we found 1120 genes to be consistently up regulated between the two affected siblings and the unaffected sibling (Table S9). Of the up regulated genes, 15 also contained allelic variant pairs (Table S3). None of the 44 differentially expressed genes with allelic variant pairs are reported in RetNet as IRD genes.

## Supplemental Data

Supplemental data includes two figures and 11 tables.

## Acknowledgments

The authors would like to thank the OGI-081 family for their participation in this study. We thank the members of the Ocular Genomics Institute Genomics Core facility for their assistance with sequencing analyses. We would like to thank Beryl Cumming for help generating the Sashimi plot in figure 2. This work was supported by grants from the National Eye Institute [R01EY012910 (EAP), R01EY026904 (KMB/EAP) and P30EY014104 (MEEI core support)], and the Foundation Fighting Blindness (EGI-GE-1218-0753-UCSD, KMB/EAP). DMG was supported by Foundation Fighting Blindness (BR-GE-1213-0632-UWI), Sandra Lemke Trout Chair in Eye Research, RRF Emmett A Humble Distinguished Directorship, McPherson Eye Research Institute, Research to Prevent Blindness. We acknowledge the Genotype-Tissue Expression (GTEx) Project that was supported by the Common Fund of the Office of the Director of the National Institutes of Health, and by NCI, NHGRI, NHLBI, NIDA, NIMH, and NINDS. We acknowledge the ENCODE Consortium and the Thomas Gingeras, CSHL production lab for the use of skin RNA-seq samples ENCSR551NII, ENCSR991HIR, ENCSR460YCS and ENCSR321PGV.

## Web Resources

ENCODE https://www.encodeproject.org/experiments/ENCSR551NII/

ExAC http://exac.broadinstitute.org/

gnomAD http://gnomad.broadinstitute.org/

GTEx Portal https://gtexportal.org/home/

Picard tools http://broadinstitute.github.io/picard/

RetNet http://www.sph.uth.tmc.edu/RetNet/

## Accession Numbers

The accession numbers for the RNAseq samples reported in this paper (BioProject: PRJNA564377) are:

SRA:SRR10082823 SRA:SRR10082822 SRA:SRR10082821

SRA:SRR10082828 SRA:SRR10082829 SRA:SRR10082824 SRA:SRR10082830 SRA:SRR10082827 SRA:SRR10082826 SRA:SRR10082825 SRA:SRR10082820

