## Supplemental data for "A combined RNA-seq and whole genome sequencing approach for identification of non-coding pathogenic variants in single families"

<u>Supplementary Information</u>

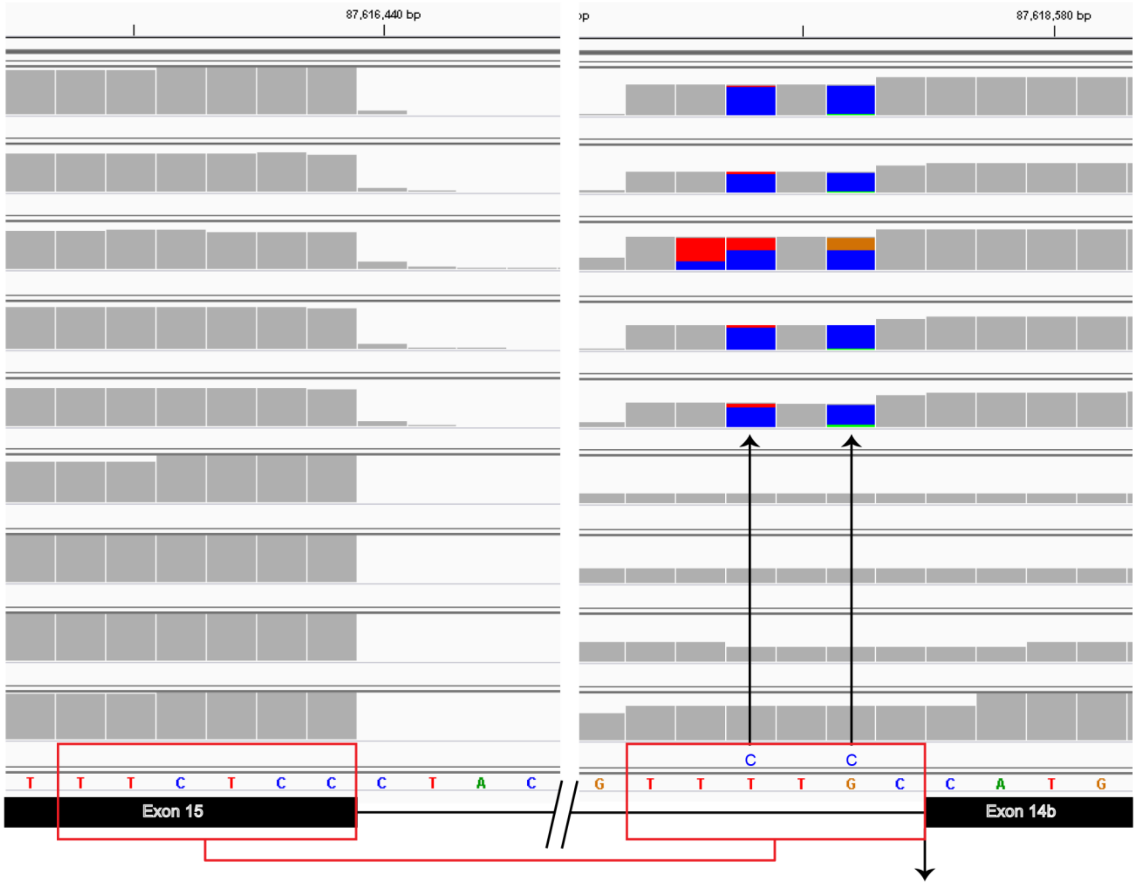


Figure S1: IGV view of the alignment error in exon 14b. IGV images of the RNA-seq reads alignment to the donor site of exon 14b and the acceptor site of exon 15. The reads spanning the splice junction are not being split and instead a six nucleotide sequence (red boxes) that belongs to exon15 is erroneously added to exon 14b causing the appearance of two mismatches (upward arrows). The correct split location is marked with a downward arrow.

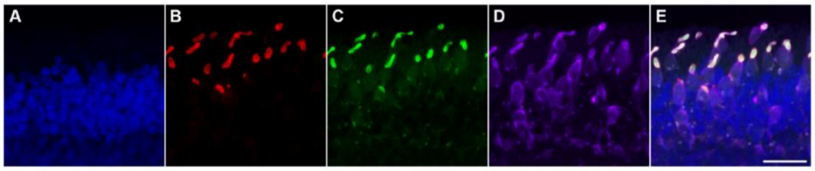


Figure S2: CNGB3 co-localizes with ML opsin in outer segments of human retina. A-E. Confocal images of adult human retina immune-stained with B. ML opsin (red), C. CNGB3 (green), D. ARR3 (purple) and E. merge with nuclei counterstained with DAPI (blue, also in A) demonstrating co-localization of CNGB3, ML opsin and ARR3 (white) in the outer segments of cones. Scale bar = 25 microns.

Table S1: Primary antibodies

| | Host | Source | Dilution |
|---|---|---|---|
| ARR3 | goat | Novus | 1:100 |
| CNGB3 | mouse | Santa Cruz Biotechnology | 1:100 |
| ML opsin | rabbit | Millipore | 1:500 |
| NR2E3 | mouse | abcam | 1:300 |
| S opsin | rabbit | Millipore | 1:500 |

Table S2: Sequin spike in controls analysis by Anaquin.

| SampleName | Intron-Exon Sensitivity (%) | IS Sensitivity (%) |
|---|---|---|
| OGI-081-197-1 | 0.91 | 0.92 |
| OGI-081-197-2 | 0.88 | 0.90 |
| OGI-081-197-3 | 0.93 | 0.94 |
| OGI-081-198-1 | 0.93 | 0.93 |
| OGI-081-198-2 | 0.91 | 0.93 |
| OGI-081-198-3 | 0.93 | 0.93 |
| OGI-081-340-1 | 0.93 | 0.93 |
| OGI-081-340-2 | 0.94 | 0.95 |
| OGI-081-340-3 | 0.91 | 0.91 |
| OGI-081-340-4 | 0.92 | 0.94 |
| OGI-081-340-5 | 0.91 | 0.92 |
